# Host-Microbiome Associations in Saliva Predict COVID-19 Severity

**DOI:** 10.1101/2023.05.02.539155

**Authors:** Hend Alqedari, Khaled Altabtbaei, Josh L. Espinoza, Saadoun Bin-Hasan, Mohammad Alghounaim, Abdullah Alawady, Abdullah Altabtabae, Sarah AlJamaan, Sriraman Devarajan, Tahreer AlShammari, Mohammed Ben Eid, Michele Matsuoka, Hyesun Jang, Christopher L. Dupont, Marcelo Freire

**Affiliations:** Department of Oral Health Policy and Epidemiology, Harvard School of Dental Medicine, Boston, MA, 02115, USA; Dasman Diabetes Institute, Kuwait; Dasman Diabetes Institute, 1180, Dasman, Kuwait; School of Dentistry, Faculty of Medicine and Dentistry. University of Alberta. Edmonton AB, T6G 2L7, Canada; Department of Pediatrics, Farwaniyah Hospital, Ministry of Health, Kuwait; Department of Pediatrics, Amiri Hospital, Ministry of Health, Kuwait; Department of Genomic Medicine and Infectious Diseases, J. Craig Venter Institute, La Jolla, CA 92037, USA; Division of Infectious Diseases and Global Public Health Department of Medicine, University of California San Diego, La Jolla, CA, USA

**Keywords:** COVID-19, saliva microbiome, inflammatory cytokines, machine learning, network analysis

## Abstract

Established evidence indicates that oral microbiota plays a crucial role in modulating host immune responses to viral infection. Following Severe Acute Respiratory Syndrome Coronavirus 2 – SARS-CoV-2 – there are coordinated microbiome and inflammatory responses within the mucosal and systemic compartments that are unknown. The specific roles that the oral microbiota and inflammatory cytokines play in the pathogenesis of COVID-19 are yet to be explored. We evaluated the relationships between the salivary microbiome and host parameters in different groups of COVID-19 severity based on their Oxygen requirement. Saliva and blood samples (n = 80) were collected from COVID-19 and from non-infected individuals. We characterized the oral microbiomes using 16S ribosomal RNA gene sequencing and evaluated saliva and serum cytokines using Luminex multiplex analysis. Alpha diversity of the salivary microbial community was negatively associated with COVID-19 severity. Integrated cytokine evaluations of saliva and serum showed that the oral host response was distinct from the systemic response. The hierarchical classification of COVID-19 status and respiratory severity using multiple modalities separately (i.e., microbiome, salivary cytokines, and systemic cytokines) and simultaneously (i.e., multi-modal perturbation analyses) revealed that the microbiome perturbation analysis was the most informative for predicting COVID-19 status and severity, followed by the multi-modal. Our findings suggest that oral microbiome and salivary cytokines may be predictive of COVID-19 status and severity, whereas atypical local mucosal immune suppression and systemic hyperinflammation provide new cues to understand the pathogenesis in immunologically naïve populations.

**Significance Statement:** The oral mucosa is one of the first sites encountered by bacterial and viral infections, including SARS-CoV-2. It consists of a primary barrier occupied by a commensal oral microbiome. The primary function of this barrier is to modulate immunity and provide protection against invading infection. The occupying commensal microbiome is an essential component that influences the immune system’s function and homeostasis. The present study showed that the host oral immune response performs unique functions in response to SARS-CoV-2 when compared to systemic responses during the acute phase. We also demonstrated that there is a link between oral microbiome diversity and COVID-19 severity. Additionally, the salivary microbiome was predictive of not only disease status but also severity.

## Introduction

The oral mucosa is one of the first sites where infections are encountered and is one of the primary barriers maintaining a low-level inflammatory response to the resident commensal microbiome(1). This barrier effectively regulates immunity and protects the body against the breakdown of defense mechanisms leading to the development of oral or systemic diseases(1). This defense mechanism involves complex interactions between the immune system, the microbiome, and the physical barriers in the oral cavity, such as saliva and mucosal tissue(2). A balanced healthy microbiome is an essential component of the host and has often been considered an organ and protective component of the human body(3), which is involved in maintaining homeostasis, epithelial cell proliferation and differentiation, and in the regulation of immune response(4). Well-established evidence showed that the immune defense of the mucosal barrier is built independently of commensal colonization and that disruption of homeostasis between the oral microorganisms leads to certain diseases(1). The interplay between the microbiome and the host forms a complex ecosystem that serves as both a metabolic and physical barrier to outcompete external microorganisms(3), such as viral invaders.

The impact of the mucosal microbiome during coronavirus disease 2019 (COVID-19) is an emerging area with ongoing research. Recent evidence suggests that the severity of COVID-19 was related to microbiome changes, as SARS-CoV-2 replicates in cells expressing angiotensin-converting enzyme 2 (ACE2) and transmembrane protease serines 2 (TMPRSS2) receptors, both of which are found in the oral and nasal mucosa(4). These ongoing investigations are starting to reveal the relationship between the oral microbiome and disease pathogenesis. The decrease in oral microbial diversity was correlated to COVID-19 patients in ICU(5), while others finding that no changes in the nasal microbiome in mildly symptomatic COVID-19-positive individuals compared to controls(6). Although evidence is emerging, there are no molecular cues with predictive potential to understand symptoms from SARS-CoV-2 and risk to severity. While systemic immune responses can be monitored through blood sampling, it does not depict the mucosal responses from a host-microbial perspective(7). To date, there is a paucity of evidence regarding the immune responses (present in fluids such as saliva and plasma) in subjects that experienced and resolved from SARS-CoV-2 infection. It is consequently important to document molecular signatures in COVID-19 patients at the mucosal level to better define normal and pathogenic convalescence processes and to detect the possible initiation of aberrant innate immune activation, especially in fluids that are in direct contact with SARS-CoV-2. Although systemic immune responses can be monitored through blood sampling, understanding oral immune competence is complex(8).

Here, in a case-control study, we measured and compared SARS-CoV-2 oral mucosal responses in matched saliva and plasma samples and investigated how the host-microbiome in saliva can predict COVID-19 status and severity. We characterized host functional cytokines impaired by SARS-CoV-2 infection by comparing responses in acute COVID-19 subjects to healthy controls and by comparing responses in saliva to plasma. This case-control study aims to evaluate whether host-microbiome in saliva can predict COVID-19 status in addition to severity.

## Results

As body fluids are exposed to different antigens, we investigated how the body responds to SARS-CoV-2 in saliva compared to those in matched serum. Paired saliva and serum samples from COVID-19 donors in the acute phase were collected and subjected to comparative analyses among demographic factors (age, gender, and initial COVID-19 disease severity) (Table 1).

**Table 1.**
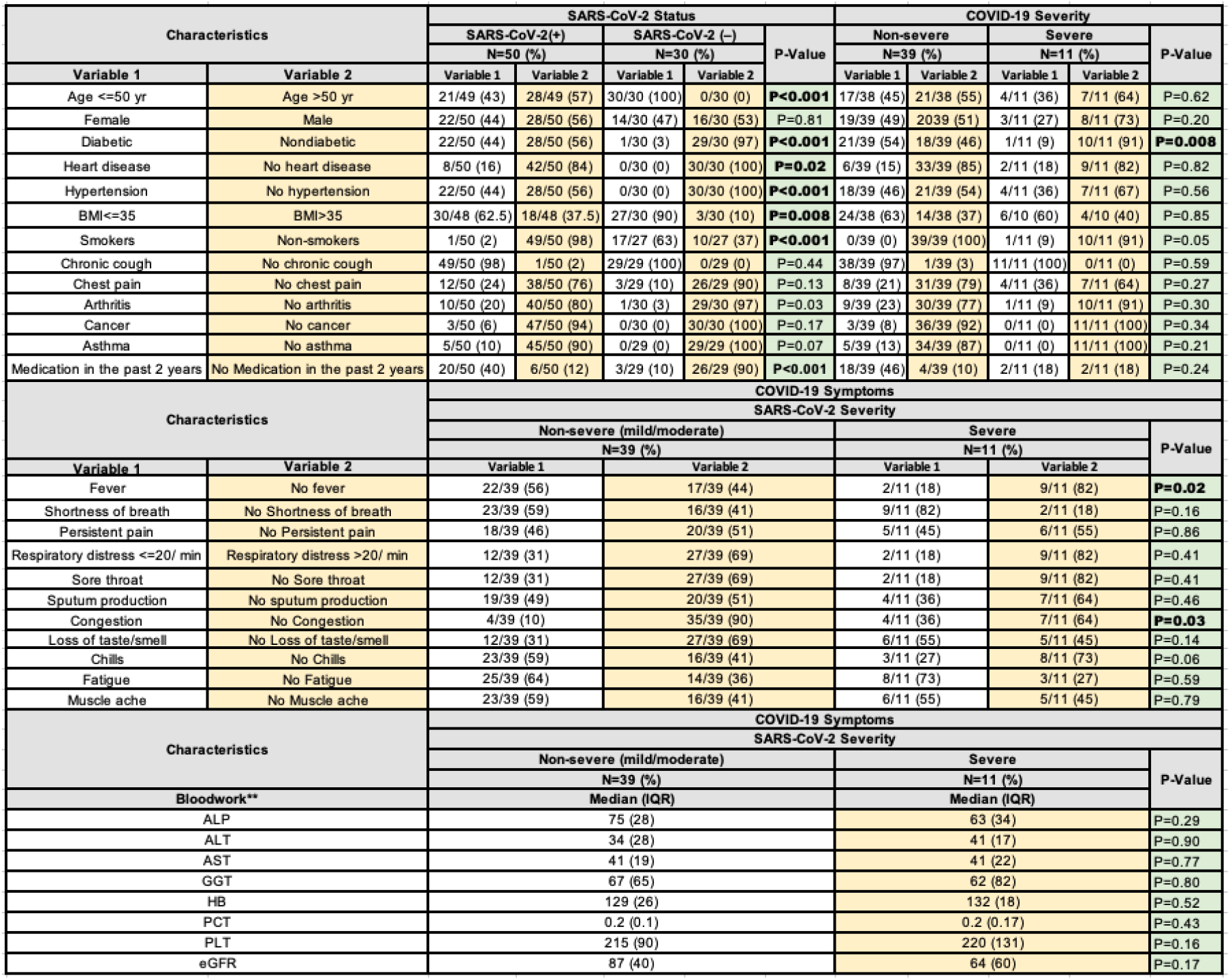
Clinical characteristics and demographics of the study patients according to disease status and severity. The chi-square test was conducted for categorical variables, and the Mann-Whitney test was conducted for continuous variables. IQR: Interquartile Range. The severity of the disease was stratified into mild: hospitalized, no oxygen therapy (n=11); moderate: hospitalized, low-flow oxygen (<10 L/min); and severe: hospitalized, high-flow oxygen (>10 L/min) (n=39). Mild and moderate groups were combined under “non-severe”. Blood work**: ALP: Alkaline phosphatase (liver function); ALT: alanine transaminase (liver function); AST: aspartate transferees (liver function); GGT: gamma-glutamyl transferase (liver function); HB: hemoglobin; PCT: procalcitonin; PLT: platelet count; eGFR: the measure of kidney function (the lower, the worse). Reference for respiratory distress cut off is 20/minute(42).

A total of 80 individuals were included in the study, consisting of 50 SARS-CoV-2 positive individuals confirmed by RT-PCR from nasopharyngeal swabs and 30 non-infected individuals. The severity of the disease was categorized as mild (hospitalized without oxygen therapy, n=11), moderate (hospitalized with low-flow oxygen <10 L/min, n=28), and severe (hospitalized with high-flow oxygen >10 L/min, n=11). The analysis combined the mild and moderate groups into a “non-severe” category. Of the participants, 55% were males and 45% were females.

Among the SARS-CoV-2 positive group, 56% were older than 50, and 64% were older than 50, among the severe cases. Diabetes was present in 44% of the SARS-CoV-2 positive group, while only 9% of the subjects with severe symptoms had diabetes. Hypertension was present in 44% of SARS-CoV-2-positive individuals and 36% of those with severe symptoms. Obesity was observed in 37% of SARS-CoV-2-positive individuals and 40% of those with severe symptoms. All subjects with severe symptoms had a chronic cough.

Respiratory distress was reported by 82% of the SARS-CoV-2 positive group, while shortness of breath was reported by 82% of the severe cases at the time of data collection. Additionally, 45% of subjects with severe symptoms experienced persistent chest pain, 36% had sputum production, and 55% had a loss of taste and smell. Only 18% of subjects with severe symptoms had fever.

The demographic variables are presented in table 1.

### Microbial composition stratified samples by COVID-19 diagnosis better than cytokines

We first evaluated the microbiome composition detected in saliva to elucidate the effect of disease on the microbiome. First, we conducted a principal coordinates analysis on Jaccard dissimilarity, which showed distinct clusters between control, mild/moderate cases, and severe cases along the first PC-axis, indicating that COVID-19 affects the microbiome composition during the acute phase of infection (p<0.05, ADONIS on Jaccard distances with FDR correction, figure 1A). To understand what drives this microbial-enhanced separation between the different conditions, we examined the alpha diversity. The control group had statistically higher diversity compared to the mild/moderate group, which in turn had statistically higher diversity than the severe group (p<0.05, observed features after FDR correction, figure 1B, supplementary figure S1). The abundance-aware Shannon diversity demonstrated that the relationship between the mild/moderate and control groups was not statistically significant (P>0.05, figure 1C) therefore, the difference in observed ASVs was driven by rarer taxa. We next conducted a principal coordinates analysis on Jaccard dissimilarity, which showed distinct clusters between control, mild/moderate cases, and severe cases along the first PC-axis, indicating that, indeed, COVID-19 affects the microbiome composition during the acute phase of infection (p<0.05, ADONIS on Jaccard distances with FDR correction, figure 1A). To examine this effect further, we broke Jaccard dissimilarity into its two components; replacement of taxa with another (turnover) and whether one sample is a subset of another (nestedness)(9). Nestedness was statistically significant between control and mild/moderate cases, but not turnover (p=0.012 and 0.299) respectively, ANOSIM after FDR correction). On the other hand, the difference between mild/moderate and severe cases was due to turnover and not nestedness (p=0.003 and 0.365). Composition-aware quantitative analysis of the microbiome also showed a similar pattern of clustering between control, mild/moderate and severe cases (p<0.05, ADONIS of PhILR distances after FDR correction, figure 1D). In addition, alpha rarefaction curves demonstrated that sufficient sample sequencing was achieved therefore the effect is not due to insufficient sequencing depth (supplementary figure S2).

**Figure 1.**
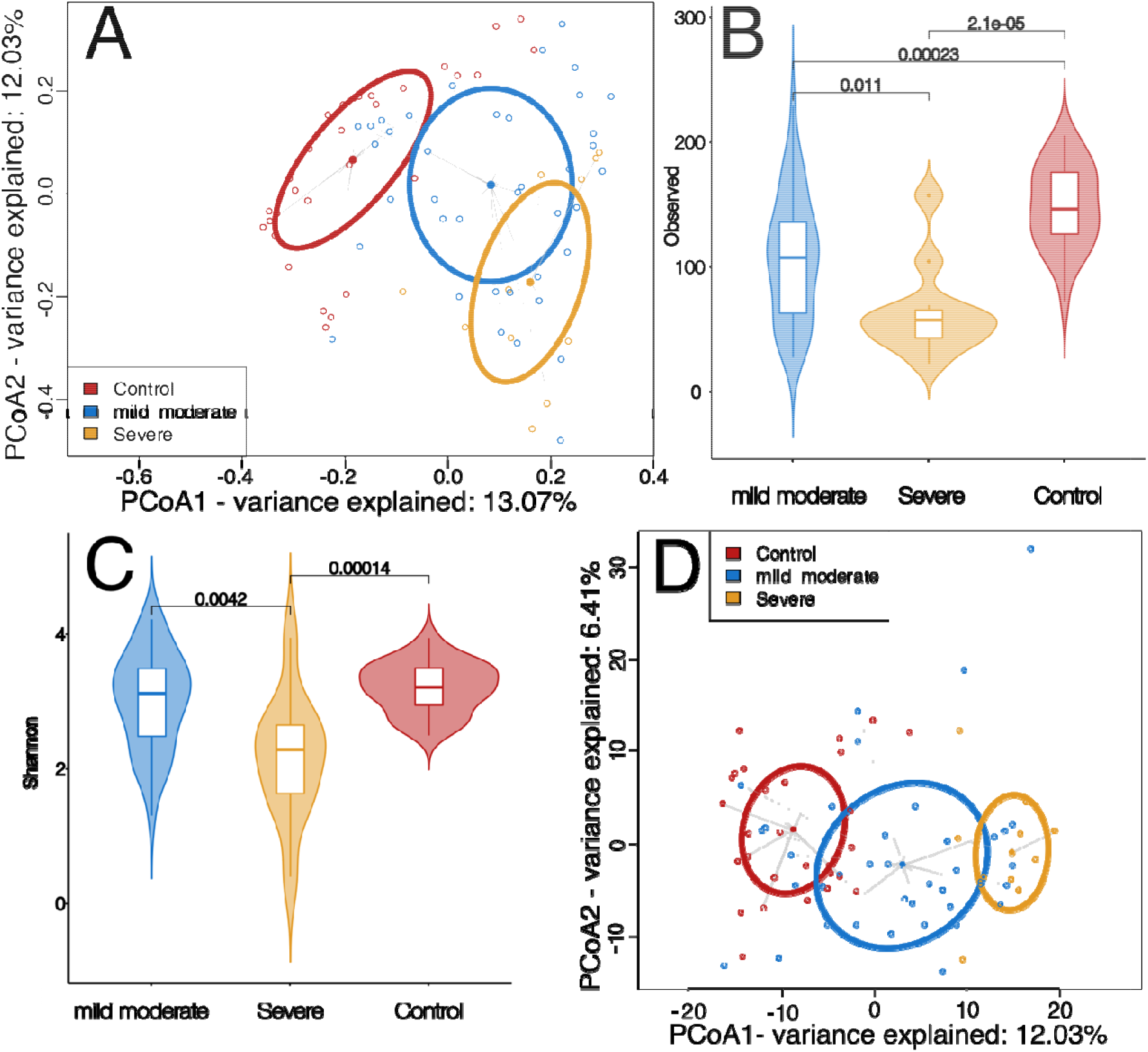
Analysis of salivary microbiome between the 3 groups, control (36), mild/moderate (blue) and severe (yellow). A: Principal coordinates analysis (PCoA) of Jaccard dissimilarity demonstrating the three clusters formed by the microbial constituents. Ellipsoids show the deviation of spread of each group (p<0.05, ADONIS). B. Violin plots of observed features diversity between the three groups (p<0.05, Wilcoxon rank-sum test). C.Violin plots of Shannon diversity between the three groups (p<0.05, Wilcoxon rank-sum test). D. Principal coordinates analysis (PCoA) of Euclidean distances of PhILR distances, demonstrating the three clusters between the conditions.

To gain insight into the capability of the oral variables in distinguishing COVID-19 status, we used unsupervised clustering (Ward hierarchical clustering) on the samples using their salivary microbiota and the salivary cytokines. The results revealed that the salivary microbiota clustered most of the COVID-19-positive patients along the first tree bifurcation, while the salivary cytokines were not able to show the same distinction (Supplementary figure S3). This finding was reflected in the predictive accuracy of each variable, where the microbiota outperformed the cytokines (Supplementary figure S3).

### Blood and saliva cytokines reflect different host responses to COVID-19

We profiled saliva and serum samples to characterize oral mucosal and systemic responses following SARS-CoV-2 infection more comprehensively. Out of a 65-cytokine assay, 15 salivary and 45 blood cytokines were significantly different between SARS-CoV-2 positive and negative patients (p<0.05, Kruskal-Walli’s test, figure 2). Only 9 of these cytokines were shared between the two profiles Furthermore, the salivary Hepatocyte Growth Factor (HGF) and Fibroblast Growth Factor 2 (FGF2) were statistically significantly lower in COVID-19-positive patients compared to the controls (<0.05), but not in blood. Additionally, salivary TNF-beta (Tumor Necrosis Factor beta), IL10 (Interleukin 10), MCP2/CCL8 (Monocyte Chemotactic Protein-2/C-C motif chemokine ligand 8), ENA78/CXCL5 (Epithelial cell-derived neutrophil-activating peptide 78/C-X-C motif chemokine ligand 5), CD40L (CD40 Ligand), IL2R (Interleukin 2 Receptor), IL12p70 (Interleukin 12), and HGF (Hepatocyte Growth Factor) were found to be expressed less in the mild/moderate condition compared to either severe or control (Supplementary figures S4-5), possibly indicating an inverse relationship where the mucosal cytokine expression is suppressed. We then determined the influence of the immune subclusters in relation to each fluid by investigating correlations. To confirm this, we examined the relationship between blood and salivary cytokines in each condition, which confirmed that while the relationship in non-SARS-CoV-2 infected individuals was positive, the inverse relationship between salivary and blood cytokines is present in severe cases (Figure 2 C-F).

**Figure 2.**
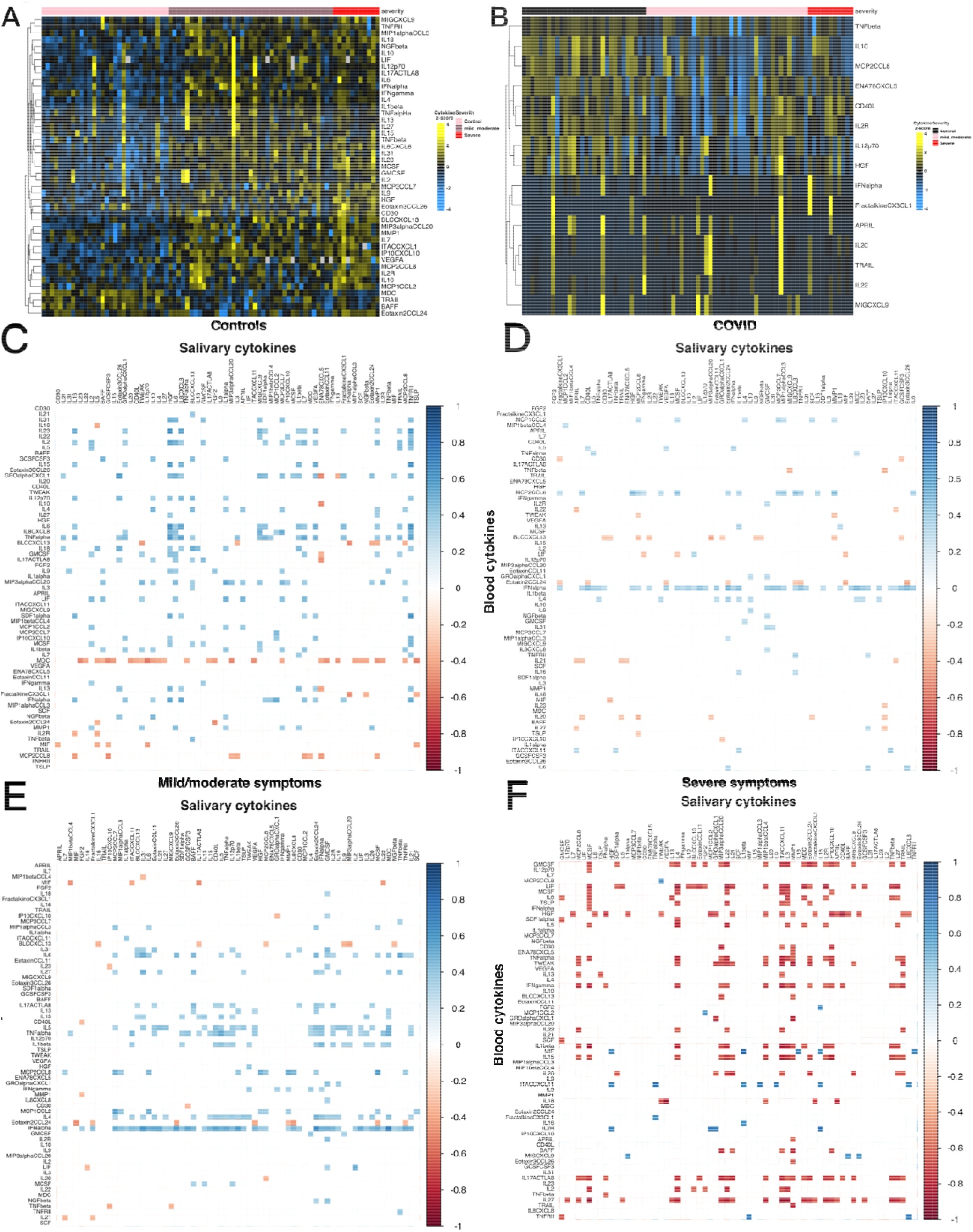
Salivary and blood cytokines. Heatmaps of statistically significant (Kruskal-Wallis) cytokines in control, mild/moderate and severe measured in A. Blood, and B. Saliva. Correlogram of C. Controls. D. All COVID patients. E. Mild/moderate category. F. Severe category, between blood (rows) and salivary (columns) cytokines. Areas coloured based on the Spearman correlation Rho and any correlations not statistically significant (P>0.05) were removed and replaced with a white square.

Our results indicate that measuring only serum levels during the COVID-19 convalescent phase does not provide a full picture of the host response mounted after SARS-CoV-2 infection. Indeed, while we confirmed that our acute COVID-19 subjects had mounted a systemic response in blood, our salivary sequencing analysis revealed novel immune signatures and altered microbiome in disease when compared to healthy controls. Overall, we demonstrated that population-based investigations of saliva can be used to map global host-microbial responses to local mucosa and begin to integrate to systemic functions.

### Microbial perturbation and cytokines serve as a robust classifier for COVID-19

We next investigated the feature selection algorithm (Clairvoyance) in a hierarchical manner (ref: https://pubmed.ncbi.nlm.nih.gov/33780444/) (Figure 3A) to better understand the strength of the signal that the microbial perturbation and cytokine fluctuations are serving as a predictive/diagnostic tool for SARS-CoV-2. We examined the strength of the potential sources of salivary and blood biomarkers and their capability in four different manners: salivary microbiome only, blood cytokine only, salivary cytokine only, and multi-modal sample-specific perturbation networks (SSPN) including the three parameters stratified by COVID-19 status and severity. The feature selection results are shown in figure 3B and 3C. We demonstrated that the microbiome-only model had the highest classification capabilities, followed by the multi-modal SSPN (microbiome: 99.4%, 100%, multi-modal SSPN: 99.5%, 91.2% for submodal y1 and y2, respectively) compared to the salivary cytokine and blood cytokine alone models. The potential markers for the microbiome did not overlap between the two submodels, with y1 being represented by 10 ASVs, and y2 being represented by 6 ASVs (Table 2A, 2B). Interestingly, most nodes in the SSPN were shared across the two submodels, inferring that the interactions (edges) between these nodes are drivers to the distinction between the two submodels. None of the microbial biomarkers were shared between sub-modals y1 (N = 10 ASVs) and y2 (N = 6 ASVs) while all the other modalities shared features between sub-modals y1 and y2 (Supplementary tables S1, S2). Central to these interactions is the systemic cytokine CXCL10/IP-10.

**Figure 3.**
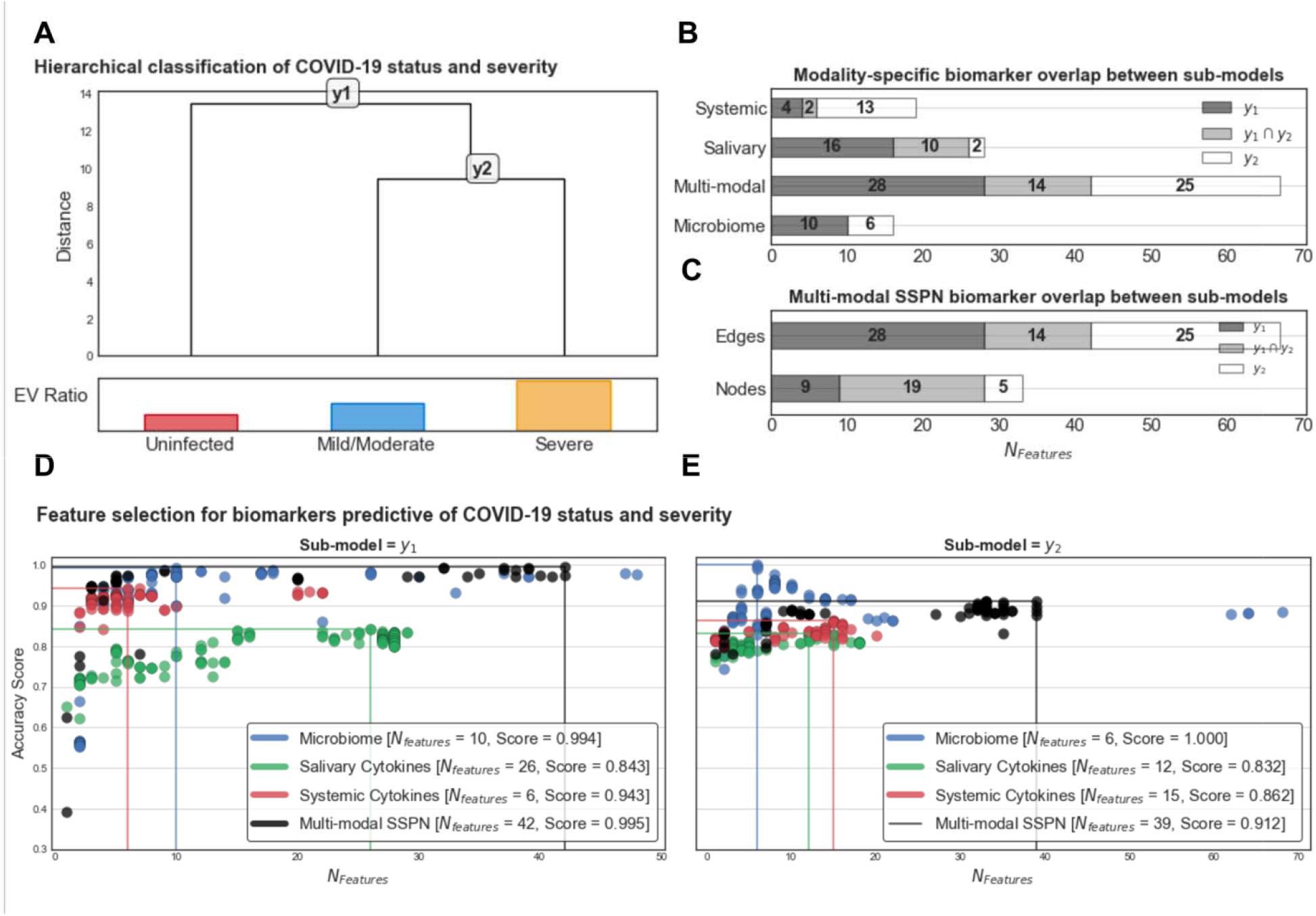
Hierarchical feature selection for identifying biomarkers predictive of COVID-19 severity. A) Hierarchical classification of COVID severity and the sub-models with relative explained variance of each group derived from 1st principal component of PCA. Modality-specific biomarker overlap between sub-models of B) abundance-based paradigms and C) inferred-interaction paradigms. In the inferred interaction paradigm, features are edges in the SSPN which can be decomposed into nodes which is not the case for the abundance-based paradigms in B. *Clairvoyance* feature selection results for D) sub-model y1 and E) sub-model y2 color coordinated by modality. Each marker on the scatter plot represents a unique hyperparameter/feature set combination that yields a specific accuracy using 10-fold cross-validation repeated with 10 different random states.

A major advantage of analyzing edge perturbations using feature selection algorithms is the ability to reconstruct edges into a network providing an opportunity to reap the benefits of graph theory. In this context, we reconstructed our aggregate networks (AN) using the multi-omic biomarkers and their fitted model weights (i.e., logistic regression coefficients) as edge weights with sub-models y1 and y2 representing ANy1 and ANy2, respectively (Figure 3). As the edge weights represent model coefficients, they can be directly interpreted as their predictive capacity in classifying stepwise severity with positive coefficients translating to a higher likelihood of a sample to be more classified as more severe and vice versa for negative coefficients.

Comparing overlap in network context can be performed at the node and edge level. For ANy1 and ANy2 there was a higher proportion of nodes shared (57.6%) compared to shared edges (20.9%) between both sub-models, indicating a common set of biomarkers useful for diagnosing COVID-19 phenotypes but the way those biomarkers interact change with severity.

In both ANy1 and ANy2, the most notable node is the systemic cytokine CXCL10/IP-10 which has disproportionately high connectivity relative to the other nodes and is highly centralized (Figure 4, S6). ANy1 and ANy2 have similar network structures but ANy1 node degree distribution is less homogenous than ANy2 with homogeneity indices of 80.7% and 88.3%, respectively. In other words, the predictive capacity of systemic cytokine CXCL10/IP-10 is disproportionately higher than the other nodes in classifying COVID-19 diagnosis (sub-model y1) relative to COVID-19 severity (sub-model y2). In sub-model y2, the classification of COVID-19 severity is more evenly distributed across the other nodes (Figure S6).

**Figure 4.**
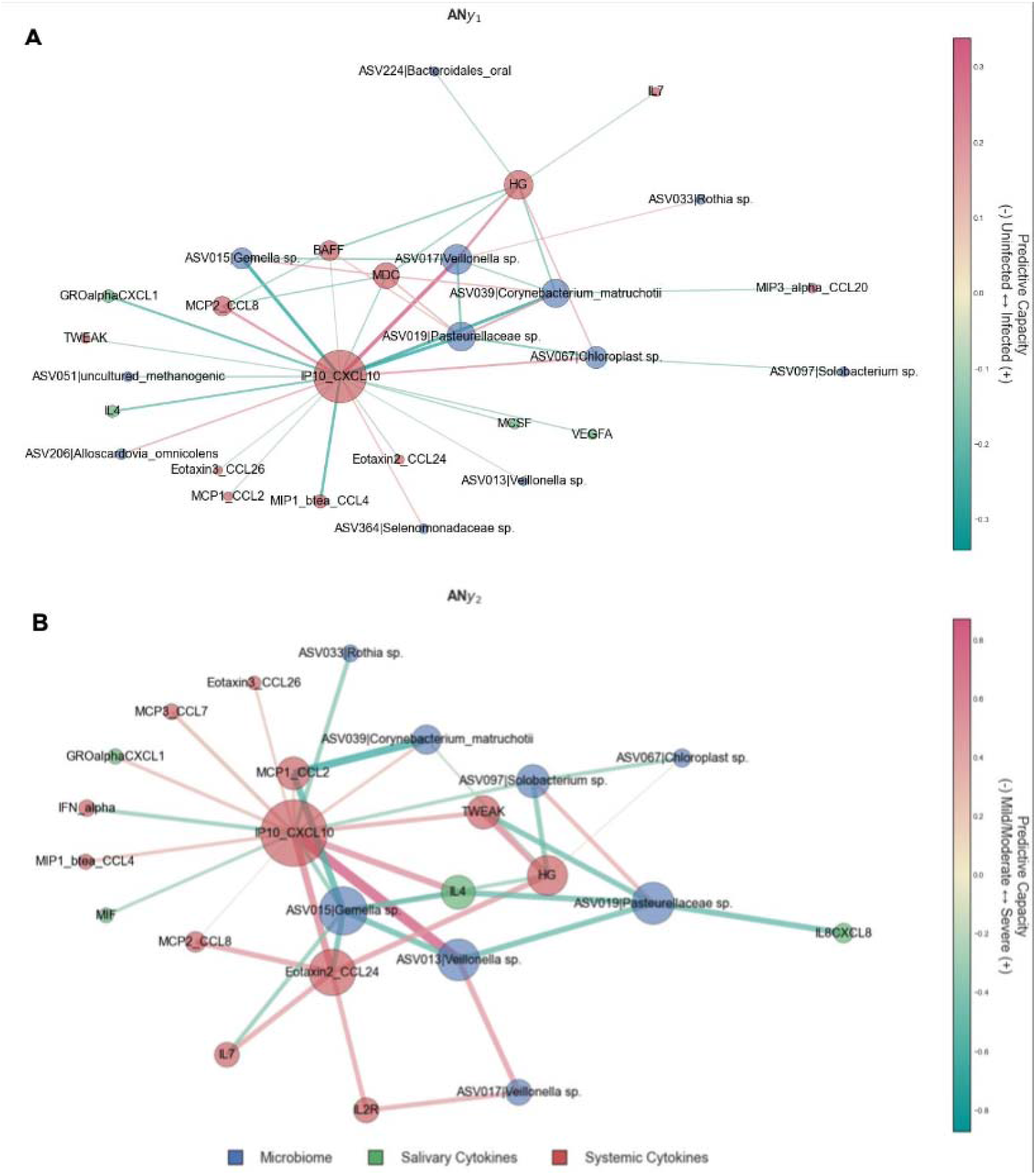
Aggregate network representations of fitted sub-models showcasing predictive capacity. – Aggregate network for sub-model A) y1 and B) y2. The edge weights can be interpreted as predictive capacity for COVID severity. For y1, positive values indicate that an increase in perturbation results in an increased likelihood that a sample is classified as y2 (i.e., infected) relative to uninfected. For y2, positive values indicate that an increase in perturbation results in an increased likelihood that a sample is classified as severe relative to mild/moderate. Node size is proportional to weighted degree as a measurement of network connectivity and, by extension, biomarker importance. C) ASVs in the SSPN as proportions of presence in the various conditions, demonstrating that the majority of the ASVs are due to differences in abundances of common taxa across all conditions, and not rare species.

For ANy1 the next highest degree nodes are ASV017 (Veillonella sp.), ASV019 (Pasteurellaceae sp.), systemic HG, and ASV039 (Corynebacterium matruchotii) and, along with CXCL10/IP-10, these 5 nodes represent 59.1% of the total predictive capacity in sub-model y1 in classifying whether a patient has been diagnosed with COVID-19 at an accuracy of 99.5% (Figure S6, S7). For ANy2 the next highest degree nodes are ASV015 (Gemella sp.), systemic CCL24|Exotaxin2, ASV019 (Pasteurellaceae sp.), ASV013 (Veillonella sp.), and systemic HG and, along with CXCL10/IP-10, these 6 nodes represent 58.4% of the total predictive capacity in sub-model y2 in classifying COVID-19 severity at an accuracy of 91.2% (Figure S6B).

## Discussion

Increasing evidence indicates that the immune system of COVID-19 subjects may be compromised when compared to healthy controls(10). Most investigations have focused on systemic body compartments such as blood and lung, and emerging investigations are dissecting the nasal mucosal responses. We demonstrated that inflammatory response to viral infection is regulated by microbiome changes locally and hypothesize that systemic responses are not predictive of systemic changes. The present analysis demonstrated that the salivary microbial community is highly susceptible to perturbation in the presence of SARS-CoV-2, leading to a distinct separation of the salivary microbiome between healthy and COVID-19-infected individuals. We also showed that the salivary microbiome was distinct between individuals with severe and mild/moderate symptoms. Our findings also demonstrate that abnormal inflammatory responses can be identified in both saliva of COVID-19 subjects, which was shown to be severity specific. This indicates that even when SARS-CoV-2 systemic responses are mounted, the local immune response does not necessarily define disease resolution. Here, we highlight saliva as an important and accessible fluid that can be monitored to identify not just antibody responses, but also diverse host-microbial pathways, including mucosal immunity, and innate immune responses.

The variation of the microbial membership detected in individuals with severe and mild/moderate symptoms can be attributed to the replacement of taxa with other taxa (turnover). However, the separation between groups (health versus disease) was mainly attributed to different phenomena; we found a decrease in the number of salivary bacterial members in the mild/moderate cases compared to the control cases, with the taxa of mild/moderate being a subset of the control. This decrease was not seen when adjusting for the overall abundance of the microbiome (Shannon diversity). Therefore, our findings suggest that the transition from control to mild/moderate disease status preserves most of the microbial membership and is mainly driven by members with higher abundance; that is, the loss of microbial membership in the mild/moderate cases occurred in less abundant taxa. This demonstrates that the salivary microbiome is generally well-adapted to perturbations, where these bacteria can maintain the overall community structure despite the change in the mucosal inflammatory environment during the acute phase of infection. Conversely, the transition from mild/moderate to severe disease corresponds to a larger environmental stress, causing a further collapse in microbial richness, thus allowing for the replacement of some taxa with others (turnover). However, when accounting for the overall collapse in membership and richness, this replacement of taxa was not enough to reverse the trend of reduced alpha diversity in severe cases.

Carefully designed immune studies aimed at implementing cytokine testing by investigations of blood-derived fluids, but not saliva. Although it is well-known that mucosal and systemic immunity responds differently to pathogens(11), this study adds to the evidence that each immunity system responds in a distinct manner to SARS-CoV-2 locally. Our findings showed that the oral response was distinct from the systemic response. The present study demonstrated that systemic cytokines showed higher levels in mild/moderate cases compared to control cases, which is expected. However, salivary cytokine levels were lower in mild/moderate cases compared to control cases. Even though the wealth of evidence agrees with the fact that cytokine production increases during the disease’s status, it was suggested that reducing cytokine production can be a strategy to manage cytokine storms and other inflammatory reactions (11). Thus, oral mucosal surfaces might lower cytokines in certain situations to balance between their two opposing roles: fighting pathogens and immune surveillance.

We further demonstrated that salivary interleukin 8 responses to SARS-COV-2 could be involved in neutrophil interactions at the oral mucosal surface. Significant correlation between salivary IL8 with blood markers, such as IL23, IL2, IL15, GROalphaCXCL1, IL12p70, IL6, TNFalpha, IL18, SDF1alpha, IL13, and IFNalpha demonstrate the tight coupling required for immune surveillance where a variety of cytokines are expressed for immune-cellular recruitment and activation of innate immune cells. This relationship is not maintained during COVID-19 infections. Additionally, we showed that the oral mucosal response was not linear; for example, salivary Interferon gamma-induced protein 10 (IP-10) demonstrates lower expression in mild/moderate conditions compared to healthy and severe cases. The importance of IP10, a pro-inflammatory chemokine for T-lymphocytes, monocytes, and NK cells, has been shown in other viral infections such as HIV and HBV infections. For example, intra-organ (liver) IP10 mRNA expression and the IP10 blood levels are concurrent in patients with chronic hepatitis B infections(12). While HIV disease progression is positively correlated with levels of IP10 in circulation(13). the positive relationship between infection level and IP10 was not found in our study, thus indicating a different response path to infection. Understanding the effects of these cytokines helps us understand the host response to the infection. IL-10 and CD40L are responsible for IgA class switching of immunoglobulins(14, 15). salivary IL12 was also reduced in mild/moderate cases compared to the severe group. IL12, a critical cytokine for the activation of dendritic cells, B-Cells and macrophages, stimulates the differentiation of T cells into Th1 cells. IL-12 also promotes the production of interferon-gamma (IFN-γ).

Taken together, the salivary cytokine profile of mild/moderate individuals supports the notion that the infection has not switched to a large-scale adaptive immune response to the virus. Further, it was suggested that the innate immune system in the oral cavity acts as the first line of defense against invading microorganisms(16), and several components of the innate immune system are involved in oral immunity, including saliva, oral epithelial cells, and other oral immune cells(16). Saliva acts as a component of innate immunity with antimicrobial activity(1), and oral epithelial cells are actively implicated in immune regulation, promote defensive immune responses, and create a constitutive tolerogenic environment that maintains immune homeostasis(17).

The relationship between the oral cavity and systemic conditions is often neglected, despite ample evidence of its connection to a multitude of diseases in distant organs, such as pneumonia(18), low birth weight(19), obesity(20), diabetes(21), and coronary disease(22). The pharynx structurally bridges the oral and nasal cavities, both of which express the ACE2/TMPRRS2 receptors that play a fundamental role in the SARS-CoV-2’s ability to infect these niches. In addition, it also contains pharyngeal tonsil tissues, an instrumental induction site for oral mucosal IgA response to respiratory viral infection. The salivary gland also plays a major role as an effector site to produce T-cell-dependent antigen-specific oral IgA response. ACE2 receptors on highly differentiated epithelial cells are also strongly associated with cytokine levels in these tissues(23, 24). Furthermore, mouse models have demonstrated that the resident microbiota on mucosal surfaces can protect against rotavirus infections(25), and it has been suggested to impact SARS-CoV-2 infections(26). The state of the mucosal tissues, such as the oral cavity, by way of perturbation of the resident microbiome and local cytokine profile, reflects the status of the host and can be used to predict and monitor the overall response. These perturbations should be further understood to gain insight into the host response of the mucosal surfaces of the upper respiratory tract.

Future studies would benefit from requiring convalescent COVID-19 subjects to report possible chronic symptoms longitudinally. The use of saliva for health monitoring has received significant attention due to its ease and simplicity of use. This study sought to explore this concept in relation to the extreme case of an exogenous infection that has recently jumped into immunologically naive humans, making it the ideal scenario for such examinations. This is particularly relevant given the evidence that depending on CT values produced by RT-PCR reflect the host’s capability to clear the virus and not the infectiousness potential of the shedding particles themselves(26, 27, 28). By closely studying this unique case, the potential of saliva monitoring as a diagnostic tool can be further understood and improved upon.

In summary, our study has found the salivary microbiome to be an excellent source to predict COVID-19 status and severity and that the oral microbiome plays an important role in the immune response by stimulating or suppressing the immune system. Our study supports the notion that the salivary microbiome could be used as a potential diagnostic tool for COVID-19 and predict severity(29). While these studies are promising, more research is needed to determine the clinical utility of using the salivary microbiome as a diagnostic tool for COVID-19. However, the potential for a non-invasive, rapid, and convenient diagnostic test using salivary microbiome analysis is exciting and warrants further investigation.

### Limitations

This study had a small sample size and may limit our ability to detect the significant distinction between different groups of symptom severity. Our COVID-19 cohort included only symptomatic subjects with mild, moderate, and severe symptoms. Asymptomatic individuals with COVID-19 may exhibit different microbiome characteristics and might be included unintentionally in the control uninfected group. In addition, the salivary microbiome can be influenced by periodontal diseases that we did not control for in the present study. Our study had limitations and future studies are needed to continue this work. While our samples were collected longitudinally, this analysis was performed for the first time point to depict the acute phase. Microbiome and inflammatory responses were not adjusted by the different baselines of each individual intervisibility. While study subjects were able to report how many days, they manifested acute COVID-19 illness (asymptomatic, mild, or moderate/severe), clinical recovery was complicated by the use of national protocols and medications such as antibiotics and immunosuppressives.

It should be noted that the microbiome results may reflect the population’s ethnicity and country of residence(30). thereby are not generalizable to all populations. To further explore this, a multi-modal analysis was conducted to identify biomarkers that could distinguish between health/disease and disease severity, with the expectation that the host response to an external infection, as in COVID-19, should be mostly conserved across different populations, with differences related to quantity, rather than activation of completely different pathways. While this study offers preliminary biomarker results, it is important to consider that the salivary microbiome is only one of many factors that could influence the course of COVID-19 and further research, such as longitudinal studies, is needed to fully understand the impact of other factors such as age, underlying health conditions, and genetic susceptibility(31).

## Materials and Methods

This study was a collaborative joint study between the Harvard School of Dental Medicine (HSDM), J Craig Venter Institute (JCVI), the Ministry of Health in Kuwait, and the University of Alberta. The study was approved by JCVI, Harvard, the Kuwait Ministry of Health, and the University of Alberta. Informed consent was obtained from all enrolled participants (Kuwait Ministry of health ethics approval: #2020/1462 Harvard: IRB21-1492, University of Alberta: Pro00125245, JVCI: exempt due to secondary analysis of de-identified samples).

### Study Design

A convenient sample strategy was used to recruit patients admitted at multiple Covid-19 centers in Kuwait between July 24th and September 4^th^, 2020. The data collection took place in multiple hospital sites at three hospitals in Kuwait: AlFarwaniyah Hospital, Jaber Al Ahmed Hospital, and Kuwait Field Hospital. Data were collected from those who provided positive consent and who tested positive for SARS-CoV-2 by RT-PCR from nasopharyngeal swabs (n=50). Non-infected individuals were employees at these hospitals who were not in contact with any COVID-19 cases (n=30). The hospital subjected them to a daily visual triage (temperature and symptoms check). However, they did not have a negative PCR test. Saliva and serum samples were collected from individuals within 48 hours of PCR-confirmed COVID-19 diagnosis. The basic demographic and clinical information (including medical history, medication, periodontal health data, sleep data, weight, height, waist circumference, neck circumference, blood group, respiratory rate, and oxygen supplementation in liters for those on oxygen) of the study participants was obtained.

The severity of the disease was stratified into mild: hospitalized, no oxygen therapy (n=11); moderate: hospitalized, low-flow oxygen (<10 L/min) (n=28); and severe: hospitalized, high-flow oxygen (>10 L/min) (n=11). Mild and moderate groups were combined under “non-severe” in the present analysis.

### Saliva collection

15 mL plastic centrifuge tubes were prelabeled with the date and subject number. We then marked the 4 mL line of the tube. A parafilm was used to stimulate saliva.

Prior to sample collection, the saliva collection tube was placed in a cup with ice. A nurse, supervised by the research coordinator, would demonstrate how to provide saliva to the patient. Each subject was instructed to take a sip of water and rinse their mouth, swallow the water, chew the piece of parafilm, and then use their tongue to push saliva as it formed into the tube. They were then instructed to place the tube back in the cup with the ice cube while they waited for more saliva to form. The saliva was collected until it reached the line (4 mL) on the tube, considering the liquid region of the saliva sample (not the foamy regions). Once the patient finished providing the saliva sample, they notified the nurse. The nurse tightened the cap on the tube, wiped it with alcohol, placed the tube in the collection rack in the cooler with ice and discarded the other materials.

### Blood collection

Serum and plasma samples were collected using standard venipuncture techniques in 7.5 mL BD Vacutainer Serum marbled topped tubes with clot activator for serum, and plasma was collected in a 5 mL lavender top tube. Samples were collected at the hospital for all samples.

### Sample processing

The samples were transferred to the Jaber Alahmad Hospital laboratory in containers with dry ice. The laboratory technician received the samples and processed them on the same day of sample collection within no more than 3 hours. Saliva samples were centrifuged at 2000 x g for 5 min, and the supernatants were separated from the pellets and transferred into different tubes. Plasma and serum samples were allowed to sit upright in racks at room temperature for 30 min prior to centrifuging at 2000xg for 10 min at room temperature. All the samples were stored at –80 °C and were transferred from Kuwait to JCVI. The samples were placed on dry ice during shipment with a monitoring device to ensure that the samples were frozen during the transfer.

### DNA extraction, library preparation, and sequencing

DNA was extracted from samples using Qiagen’s AllPrep Bacterial DNA/RNA/Protein Kit (Cat# 47054; QIAGEN, Hilden, Germany) according to the manufacturer’s instructions. Step 3 was modified to use MP Biomedicals™ FastPrep-24™ Classic Bead Beating Grinder and Lysis System (Cat# MP116004500; FisherScientific) for 1 minute instead of vortexing for 10 minutes. All recommended products were used, and, at step 2, ß-ME was used instead of DTT. Sequencing was done on the Illumina MiSeq platform. V4 hypervariable region was sequenced using the following forward (MSV4Read1) and reverse primers (MSV34Index1), 5’-TATGGTAATTGTGTGCCAGCMGCCGCGGTAA-3’ and 5’-ATTAGAWACCCBDGTAGTCCGGCTGACTGACT-’3.

Samples were deposited in NIH SRA under the accession number PRJNA948421.

### Cytokine abundance measurements

Serum samples were analyzed using the Immune Monitoring 65-Plex Human ProcartaPlex Panel (Cat# EPX650-16500-901; ThermoFisher Scientific, Vienna, Austria), according to manufacturer’s instructions, and run on the Luminex 200 system (Luminex Corporation, Austin, Texas, USA). This kit measured immune function by analyzing 65 protein targets in a single well, including cytokine, chemokine, growth factors/regulators, and soluble receptors. The provided standard was diluted 4-fold to generate a standard curve, and high and low controls were also included.

### Amplicon sequence variants (ASVs)

Analysis was performed at the amplicon sequence variant level: DNA sequences containing no sequencing errors after algorithmic correction. The paired-end DADA2(32) workflow of QIIME2 v2022.2.0(33) was used for detecting ASVs and quantifying abundance implemented using the amplicon.py module of VEBA(34). More specifically, this workflow uses the following protocol: 1) qiime tools import of paired-end reads; 2) DADA2 denoising of paired reads and ASV detection by qiime dada2 denoise-paired (forward_trim=251, reverse_trim=231, min-overlap=12); 3) taxonomic classification of ASVs using the precompiled silva-138-99-nb-classifier.qza model (Silva_v1383 SSURef_NR9); and 4) conversion into generalized machine-readable formats (e.g., QIIME2 Artifact and BIOM formatted files → tab-separated value and Fasta-formatted files).

### Cytokine and microbiome statistical analysis

Alpha diversity (i.e., richness) was calculated by summing each sample’s detected ASVs. Beta diversity (i.e., hierarchical clustering and PCoA) was calculated by computing the pairwise Aitchison distance, center log-ratio followed by Euclidean distance, of samples and using this distance matrix as input in the Agglomerative and PrincipleCoordinateAnalysis objects of Soothsayer(35) which are wrappers around SciPy(36) and Scikit-Bio(37), respectively.

PCoA with Jaccard and PhILR(38) alpha diversity, probability density function for Jaccard breakdown (betapart package in R) were conducted via the automated pipeline FALAPhyl: https://github.com/khalidtab/falaphyl/. Cytokine Net MFI values were used for cytokine measurements. Values were transformed by log10 transformation, then converted into z-scores, and finally plotted with the R package ComplexHeatmap(39). Cytokines with negative MFI values (i.e. lower signal than standard) had the lowest value plus one added to all the cytokine measurements so that the lowest value equals one so that the values can undergo the same log10 transformation and z-score standardization above. Correlations between blood and saliva cytokines were done with Spearman correlation in R, and those found to be statistically significant were then plotted with the R package corrplot(40).

Feature selection for identifying biomarkers of COVID-19 diagnosis feature selection was performed using the Clairvoyance algorithm (v2023.1.3), an extension of the Soothsayer(35) Python package using two paradigms (i.e., abundance and inferred interactions). A hierarchical classification scheme was used where the multi-class prediction [Uninfected, Mild/Moderate, Severe] was split out into two binary classifications; that is, sub-model = y1 [Uninfected vs. y2] and sub-model = y2 [Mild/Moderate vs. Severe] (Figure 5). Feature selection for sub-models y1 and y2 identify biomarkers that are informative in discriminating between uninfected and infected and mild/moderate versus. severe phenotypes, respectively.

For the abundance paradigm, we ran feature selection models using the following transformations: 1) the quality assessed center log-ratio transformed ASV abundances (multiplicative replacement of 1); and 2) the Z-score normalized cytokine levels. The predictive features that were identified using Clairvoyance on the ASV and cytokine abundances were used to build SSPNs (described below). For the inferred interaction paradigm, the perturbation matrix that is made of all SSPNs was used as input into the final rounds of feature selection.

To avoid performing exhaustive feature selection using every combination of pairwise multi-modal associations (∼176k associations where the majority of associations would contain non-informative features), we identified biomarkers for each modality individually and used the resulting informative features (y1 + y2 feature sets for each modality) as input for the SSPN analysis to produce a perturbation matrix as implemented in Nabwera & Espinoza et al. 2021(35).

Briefly, a perturbation matrix with edge weights of the fully connected SSPNs was fed into a feature selection algorithm to determine the minimal set of edges with the highest predictive capacity. That is biomarkers for COVID-19 as multi-modal associations rather than the abundances of specific features. Once the minimal edges predictive of COVID-19 are identified, the Logistic Regression classification model is fit, and the coefficients representing predictive capacity are used as edge weights to construct an aggregate network (Fig. 3). Positive coefficients indicate that an increase in perturbation corresponds with an increased likelihood of a COVID-19 diagnosis, while negative coefficients indicate that an increase in perturbation corresponds with a decreased likelihood of a COVID-19 diagnosis. Ultimately, this procedure resulted in 8 feature selection runs (y1 and y2 classifications for each scenario) that could be run on a local machine: 1-2) ASV abundances; 3-4) Salivary cytokines abundances; 5-6) Systemic cytokines abundances; and 7-8) multi-modal perturbations (i.e., SSPN).

For Clairvoyance, we used a logistic regression classifier with the following parameter space: penalty {l1,l2} and C {1e-10,0.2,0.3,0.4,0.5,0.6,0.7,0.8,0.9,1.0} with the liblinear solver and 1000 maximum iterations for convergence. The weights from the fitted classifiers are representative of a feature’s predictive capacity, that is, the coefficient of the fitted linear model. Linear models have both positive and negative coefficients. In the case of our models, a positive coefficient for a feature indicates that an increased value of the feature corresponds with an increased likelihood of the query sample to be predicted as the following: y1) infected relative to uninfected; and y2) severe relative to mild/moderate severity. Only feature sets with less than the number of samples (NSamples = 80) were considered for downstream analysis to prevent overfitting.

### Multi-omic sample-specific perturbation networks (SSPNs)

SSPNs were created to identify which inferred interactions were perturbed by adding a query sample to a reference cohort. Cytokine Z-score normalized data was propagated from previous analysis as they are not influenced by intra-sample features, while ASV abundance was retransformed using the feature subset identified by Clairvoyance which is influenced by intra-sample features. Pearson correlation was used for both continuous and multi-modal interactions (i.e., cytokine-cytokine and cytokine-ASV), while rho proportionality was used for compositional interactions (i.e., ASV – ASV). The reference category was the uninfected class, and in the y1 scenario, we used a dummy class to obtain perturbation measurements that reflect how adding individual reference samples to the larger reference cohort perturbs the associations. SSPNs were implemented using the SampleSpecificPerturbationNetwork object in the Ensemble NetworkX Python package (v2023.2.14) (35, 41) using 1000 iterations and a sample size of 0.618 with median and median-absolute deviation as the edge reduction statistic and variability measurements, respectively.

### Multi-omic aggregate networks

The logistic regression classifier used for constructing the aggregate networks was fit using the perturbation of inferred interactions (i.e., the perturbation matrix from the SSPN analysis) with the predictive feature subset identified by Clairvoyance (i.e., multi-modal edges in the perturbation matrix) and the COVID-19 severity as classes. As each feature in our classification models represented an edge, we reformatted our fitted models into undirected aggregate networks where each node represents either an ASV or cytokine, each edge represents an inferred interaction, and edge weight represents the predictive capacity of each edge when its associations are perturbed. Salivary and systemic cytokines were kept separate during all analyses.

Node degree is calculated as the sum of absolute values of edge weights connected to a node. Node degree homogeneity is a variant of Pielou’s ecological index for evenness calculated via H/llog2(Number of nodes) where H indicates Shannon entropy (ref: https://www.sciencedirect.com/science/article/abs/pii/0022519366900130).

## Acknowledgments

We acknowledge the funding agencies for the support to this work. This study was funded by J Craig Venter Institute, CA, USA; Kuwait Ministry of Health; Dasman Diabetes Institute, Kuwait; and L’Oréal-UNESCO. MF and HJ were funded by NIH R21DE029625 and Conrad Prebys Foundation grant #53.CLD and JLE were funded by NIH 1U54GH009824 and 1R01AI170111-01 to CLD.

## References

1. Moutsopoulos NM, Konkel JE. Tissue-Specific Immunity at the Oral Mucosal Barrier. Trends Immunol. 2018;39(4):276–87.

2. Şenel S. An Overview of Physical, Microbiological and Immune Barriers of Oral Mucosa. Int J Mol Sci. 2021;22(15).

3. Salzano FA, Marino L, Salzano G, Botta RM, Cascone G, D’Agostino Fiorenza U, et al. Microbiota Composition and the Integration of Exogenous and Endogenous Signals in Reactive Nasal Inflammation. J Immunol Res. 2018;2018:2724951.

4. Zhu F, Zhong Y, Ji H, Ge R, Guo L, Song H, et al. ACE2 and TMPRSS2 in human saliva can adsorb to the oral mucosal epithelium. J Anat. 2022;240(2):398–409.

5. Haran JP, Bradley E, Zeamer AL, Cincotta L, Salive MC, Dutta P, et al. Inflammation-type dysbiosis of the oral microbiome associates with the duration of COVID-19 symptoms and long COVID. JCI Insight. 2021;6(20).

6. Gupta A, Karyakarte R, Joshi S, Das R, Jani K, Shouche Y, et al. Nasopharyngeal microbiome reveals the prevalence of opportunistic pathogens in SARS-CoV-2 infected individuals and their association with host types. Microbes Infect. 2022;24(1):104880.

7. Mehta P, McAuley DF, Brown M, Sanchez E, Tattersall RS, Manson JJ. COVID-19: consider cytokine storm syndromes and immunosuppression. Lancet. 2020;395(10229):1033–4.

8. Williams DW, Greenwell-Wild T, Brenchley L, Dutzan N, Overmiller A, Sawaya AP, et al. Human oral mucosa cell atlas reveals a stromal-neutrophil axis regulating tissue immunity. Cell. 2021;184(15):4090–104.e15.

9. Baselga A, Cdl O. Betapart: An R package for the study of beta diversity. Methods in Ecology and Evolution. 2012;808-812:808–12.

10. Mathew D, Giles JR, Baxter AE, Oldridge DA, Greenplate AR, Wu JE, et al. Deep immune profiling of COVID-19 patients reveals distinct immunotypes with therapeutic implications. Science. 2020;369(6508).

11. Russell MW, Moldoveanu Z, Ogra PL, Mestecky J. Mucosal Immunity in COVID-19: A Neglected but Critical Aspect of SARS-CoV-2 Infection. Front Immunol. 2020;11:611337.

12. Willemse SB, Jansen L, de Niet A, Sinnige MJ, Takkenberg RB, Verheij J, et al. Intrahepatic IP-10 mRNA and plasma IP-10 levels as response marker for HBeAg-positive chronic hepatitis B patients treated with peginterferon and adefovir. Antiviral Res. 2016;131:148–55.

13. Lei J, Yin X, Shang H, Jiang Y. IP-10 is highly involved in HIV infection. Cytokine. 2019;115:97–103.

14. Cerutti A, Zan H, Schaffer A, Bergsagel L, Harindranath N, Max EE, et al. CD40 ligand and appropriate cytokines induce switching to IgG, IgA, and IgE and coordinated germinal center and plasmacytoid phenotypic differentiation in a human monoclonal IgM+IgD+ B cell line. J Immunol. 1998;160(5):2145–57.

15. Malisan F, Brière F, Bridon JM, Harindranath N, Mills FC, Max EE, et al. Interleukin-10 induces immunoglobulin G isotype switch recombination in human CD40-activated naive B lymphocytes. J Exp Med. 1996;183(3):937–47.

16. Kaczor-Urbanowicz KE, Martin Carreras-Presas C, Aro K, Tu M, Garcia-Godoy F, Wong DT. Saliva diagnostics - Current views and directions. Exp Biol Med (Maywood). 2017;242(5):459–72.

17. De Maio F, Posteraro B, Ponziani FR, Cattani P, Gasbarrini A, Sanguinetti M. Nasopharyngeal Microbiota Profiling of SARS-CoV-2 Infected Patients. Biological Procedures Online. 2020;22(1):18.

18. Azarpazhooh A, Leake JL. Systematic review of the association between respiratory diseases and oral health. J Periodontol. 2006;77(9):1465–82.

19. Teshome A, Yitayeh A. Relationship between periodontal disease and preterm low birth weight: systematic review. Pan Afr Med J. 2016;24:215.

20. Godlewski AE, Veyrune JL, Nicolas E. [Obesity and oral health: risk factors of obese patients in dental practice]. Odontostomatol Trop. 2008;31(123):25–32.

21. Kudiyirickal MG, Pappachan JM. Diabetes mellitus and oral health. Endocrine. 2015;49(1):27–34.

22. Janket SJ, Baird AE, Chuang SK, Jones JA. Meta-analysis of periodontal disease and risk of coronary heart disease and stroke. Oral Surg Oral Med Oral Pathol Oral Radiol Endod. 2003;95(5):559–69.

23. Zang R, Gomez Castro MF, McCune BT, Zeng Q, Rothlauf PW, Sonnek NM, et al. TMPRSS2 and TMPRSS4 promote SARS-CoV-2 infection of human small intestinal enterocytes. Sci Immunol. 2020;5(47).

24. Stocker N, Radzikowska U, Wawrzyniak P, Tan G, Huang M, Ding M, et al. Regulation of angiotensin-converting enzyme 2 isoforms by type 2 inflammation and viral infection in human airway epithelium. Mucosal Immunol. 2023;16(1):5–16.

25. Dani N, Herbst RH, McCabe C, Green GS, Kaiser K, Head JP, et al. A cellular and spatial map of the choroid plexus across brain ventricles and ages. Cell. 2021;184(11):3056–74.e21.

26. Ngo VL, Gewirtz AT. Microbiota as a potentially-modifiable factor influencing COVID-19. Curr Opin Virol. 2021;49:21–6.

27. Han MS, Byun JH, Cho Y, Rim JH. RT-PCR for SARS-CoV-2: quantitative versus qualitative. Lancet Infect Dis. 2021;21(2):165.

28. Shah S, Singhal T, Davar N, Thakkar P. No correlation between Ct values and severity of disease or mortality in patients with COVID 19 disease. Indian J Med Microbiol. 2021;39(1):116–7.

29. Sri Santosh T, Parmar R, Anand H, Srikanth K, Saritha M. A Review of Salivary Diagnostics and Its Potential Implication in Detection of Covid-19. Cureus. 2020;12(4):e7708.

30. Mason MR, Nagaraja HN, Camerlengo T, Joshi V, Kumar PS. Deep sequencing identifies ethnicity-specific bacterial signatures in the oral microbiome. PLoS One. 2013;8(10):e77287.

31. Hu J, Wang Y. The Clinical Characteristics and Risk Factors of Severe COVID-19. Gerontology. 2021;67(3):255–66.

32. Callahan BJ, McMurdie PJ, Rosen MJ, Han AW, Johnson AJ, Holmes SP. DADA2: High-resolution sample inference from Illumina amplicon data. Nat Methods. 2016;13(7):581–3.

33. Bolyen E, Rideout JR, Dillon M, Bokulich N, Abnet C, Al-Ghalith G, et al. Reproducible, interactive, scalable and extensible microbiome data science using QIIME 2. Nature Biotechnology. 2019;37:1.

34. Espinoza JL, Dupont CL. VEBA: a modular end-to-end suite for in silico recovery, clustering, and analysis of prokaryotic, microeukaryotic, and viral genomes from metagenomes. BMC Bioinformatics. 2022;23(1):419.

35. Espinoza JL, Dupont CL, O’Rourke A, Beyhan S, Morales P, Spoering A, et al. Predicting antimicrobial mechanism-of-action from transcriptomes: A generalizable explainable artificial intelligence approach. PLoS Comput Biol. 2021;17(3):e1008857.

36. Virtanen P, Gommers R, Oliphant TE, Haberland M, Reddy T, Cournapeau D, et al. SciPy 1.0: fundamental algorithms for scientific computing in Python. Nat Methods. 2020;17(3):261–72.

37. Team TS-BD. Scikit-Bio: A Bioinformatics Library for Data Scientists, Students, and Developers. 2020.

38. Silverman JD, Washburne AD, Mukherjee S, David LA. A phylogenetic transform enhances analysis of compositional microbiota data. Elife. 2017;6.

39. Gu Z, Eils R, Schlesner M. Complex heatmaps reveal patterns and correlations in multidimensional genomic data. Bioinformatics. 2016;32(18):2847–9.

40. Wei T SV. R package ‘corrplot’: Visualization of a Correlation Matrix 2021 [Available from: https://github.com/taiyun/corrplot.

41. Hagberg AA, Schult DA, Swart P, editors. Exploring Network Structure, Dynamics, and Function using NetworkX2008.

42. Park SB KD. Tachypnea StatPearls: StatPearls Publishing; 2023 [Available from: https://www.ncbi.nlm.nih.gov/books/NBK541062/.

