## Supplementary figures and tables for "Host-Microbiome Associations in Saliva Predict COVID-19 Severity"

**Affiliations**

**This PDF file includes:**

Figures S1 to S7
Tables S1 to S2

**Supplementary Figure S1**

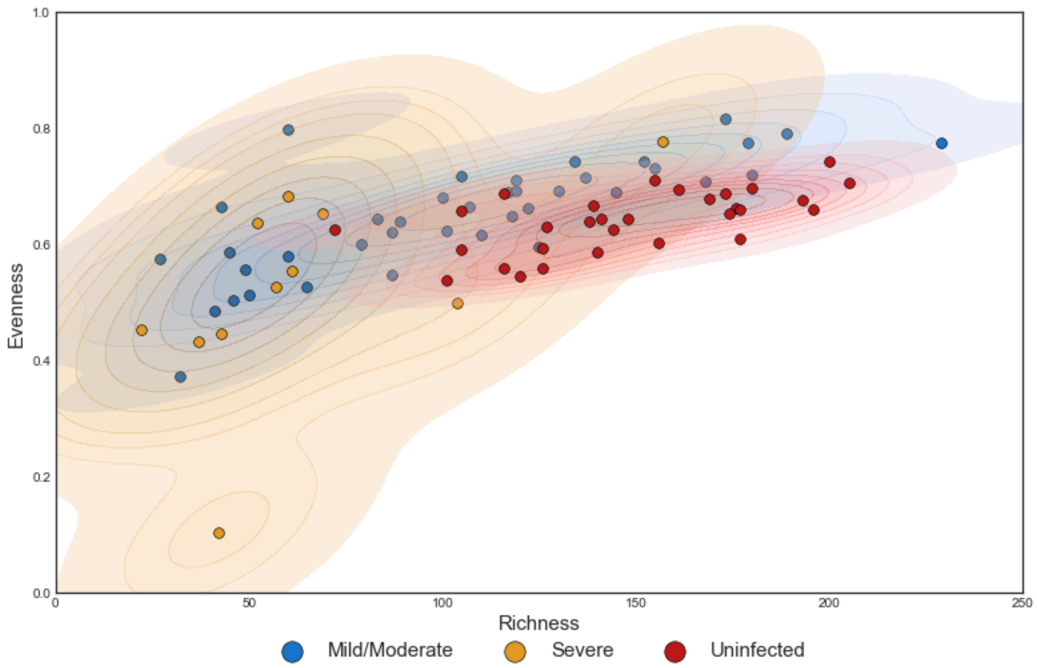


**Supplementary Figure S1 –** Scatterplots of Salivary microbiome richness (x-axis) vs. Pielou's evenness index (y-axis). Each sample is represented by a dot, and colored based on their disease status (uninfected/control – red, mild/moderate symptoms – blue, severe – yellow). Heatmap superimposed on the scatterplot to demonstrate density function.

**Supplementary Figure S2**

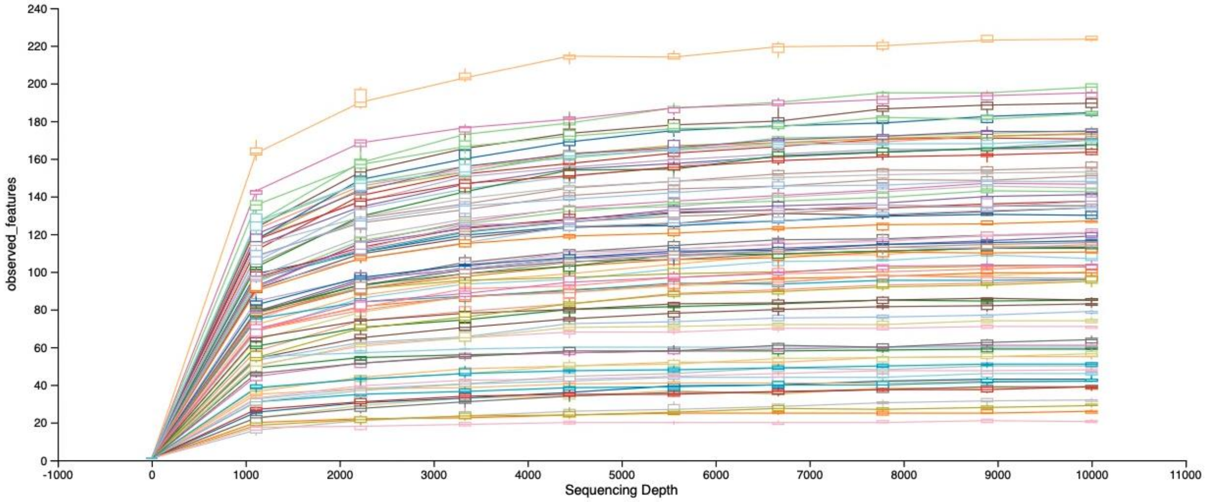


**Supplementary Figure S2.** Alpha rarefaction curves – showing “observed features” curves. Flattening of the curve demonstrates that we have achieved adequate sequencing depth for sequences.

**Supplementary Figure S3**

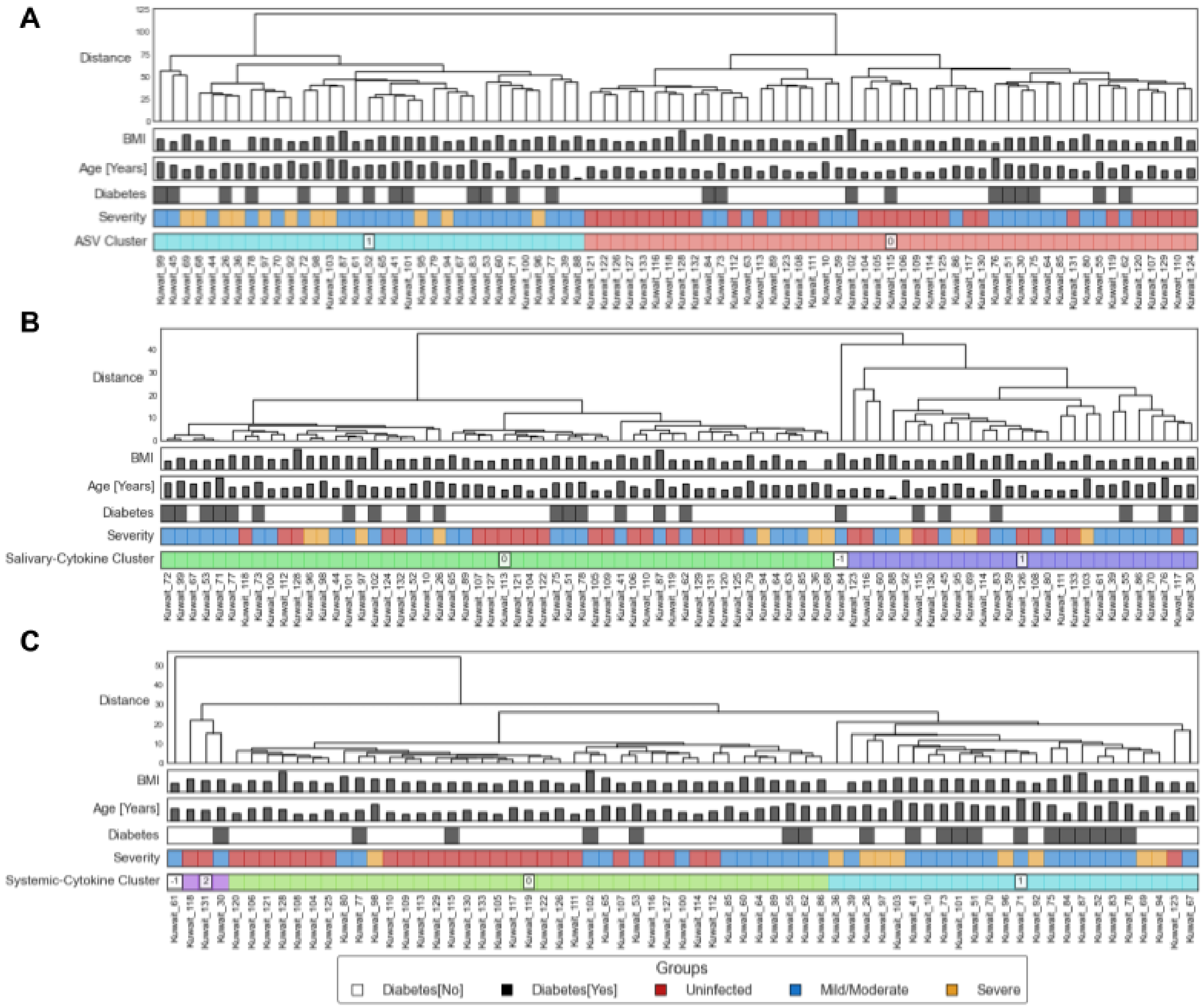


**Supplementary Figure S3.** Unsupervised clustering of saliva microbiome and cytokines. (A) Clustering using ASV abundances. (80 samples, 463 ASVs). Bottom row showing the first tree bifurcation, which has clustered most COVID-19 positive samples together. (B) Clustering using salivary cytokine abundances. (80 samples, 65 cytokines). (C) Clustering using systemic cytokine abundances. (80 samples, 65 cytokines).

### Supplementary Figure S4

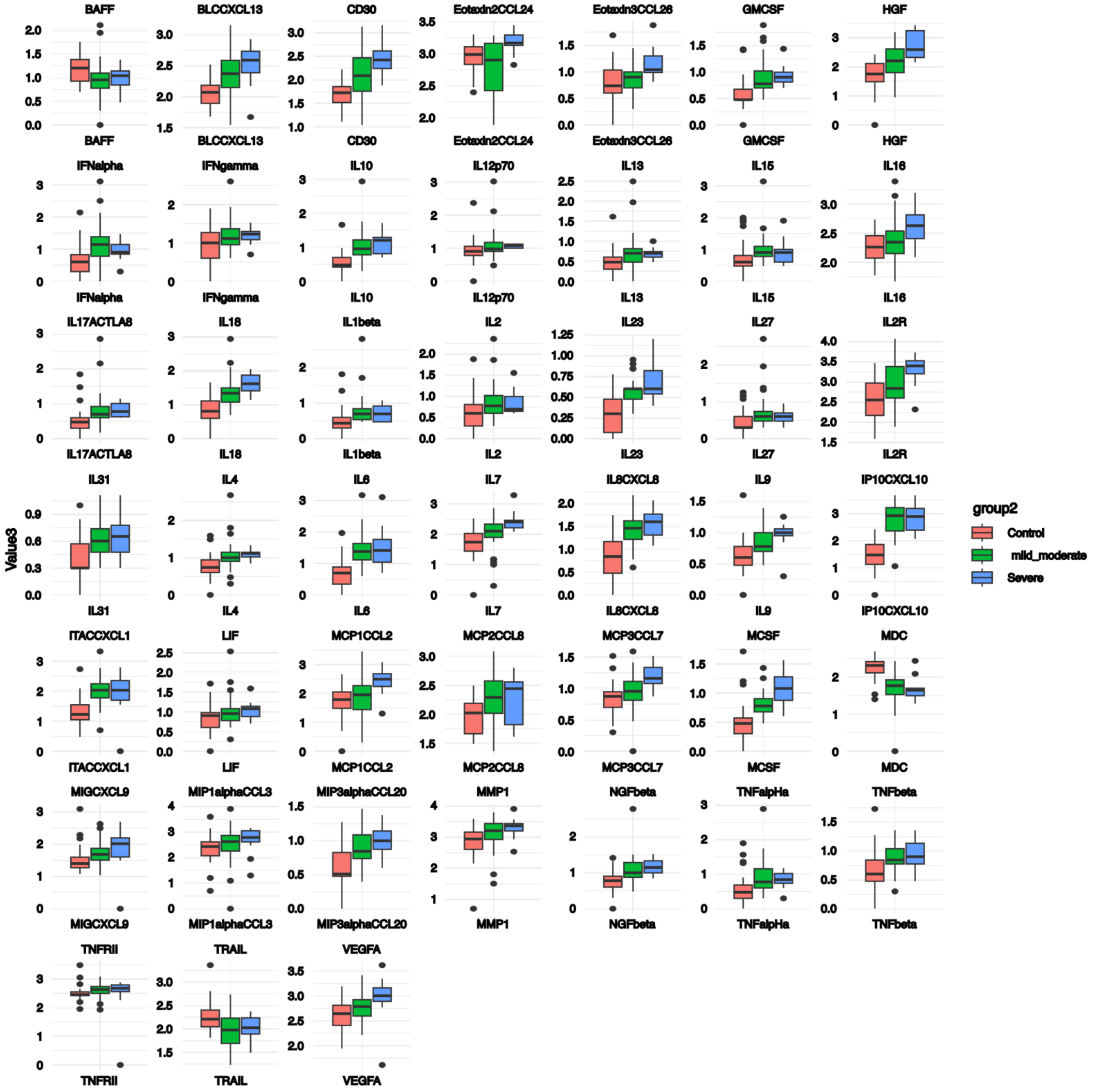


**Supplementary Figure S4.** Boxplot of representative blood cytokines ($p<0.05$, Kruskal-Walli's test). All values were log10 transformed before graphing. Whiskers demonstrate IQR.

**Supplementary Figure S5**

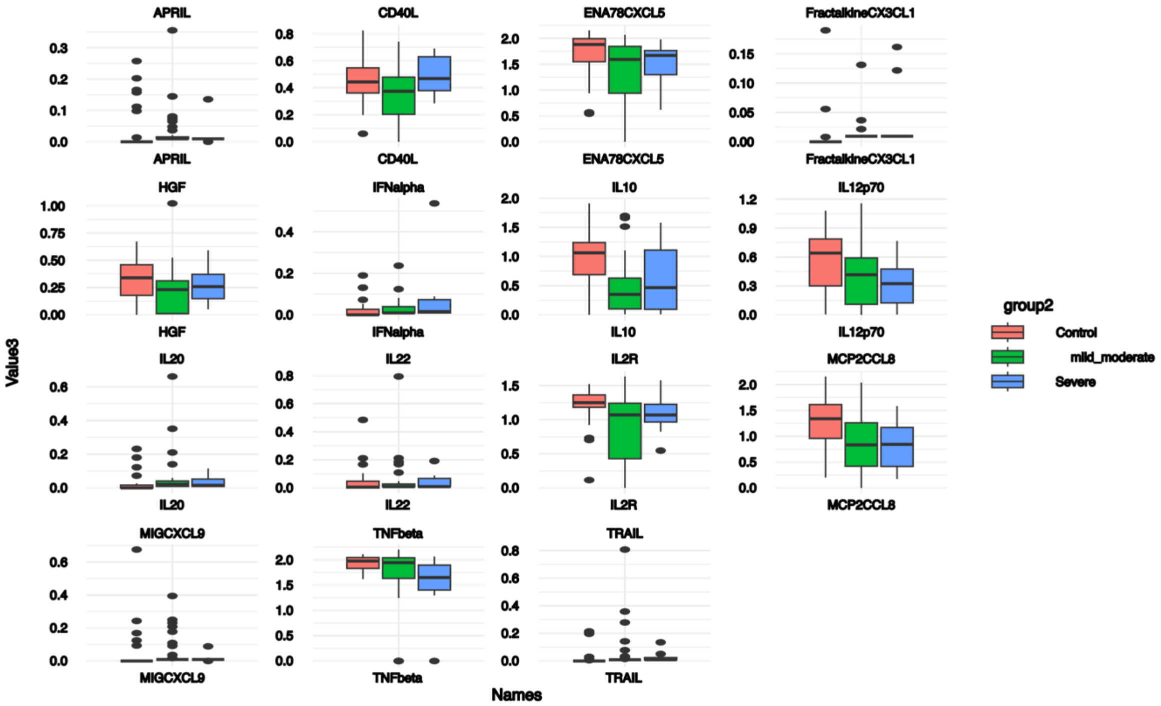


**Supplementary Figure S5.** Boxplot of representative saliva cytokines. ($p<0.05$, Kruskal-Walli's test). All values were log10 transformed before graphing. Whiskers demonstrate IQR.

**Supplementary Figure S6**

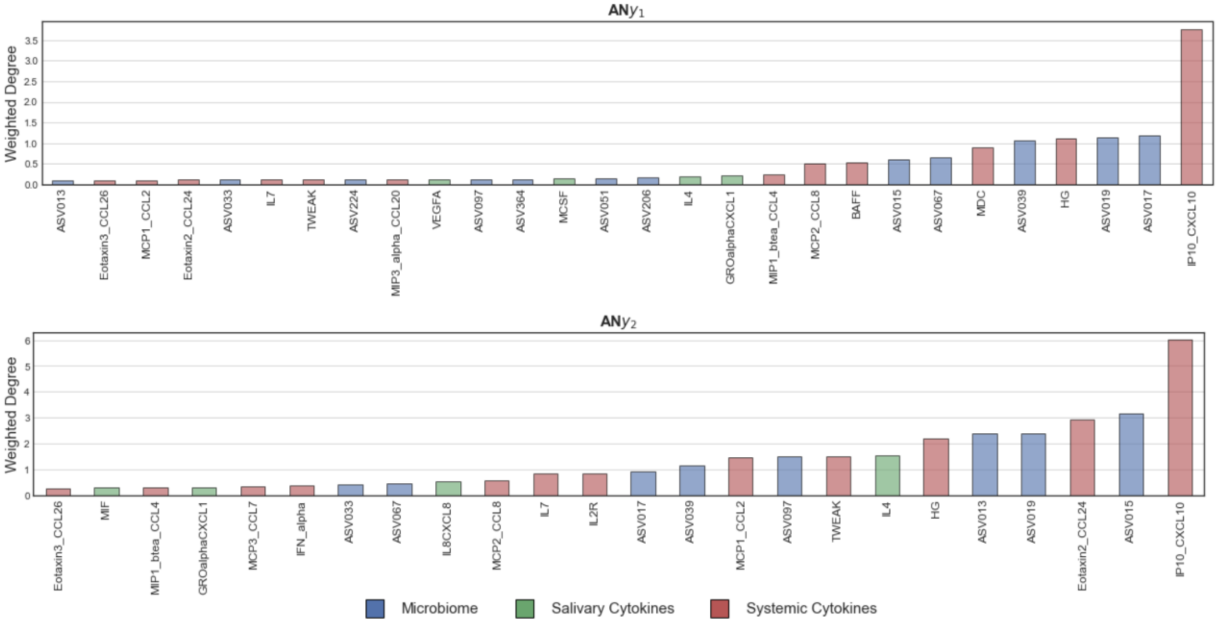


**Supplementary Figure S6**. Weighted degree of nodes in aggregate networks for sub-models y1 and y2. Node the high degree of connectivity of IP10/CXCL10 in y1 – the sum of degrees of ASV017 (Veillonella sp.), ASV019 (Pasteurellaceae sp.), systemic HG, and ASV039 (Corynebacterium matruchotii) s is 4.55 while CXCL10/IP-10 alone is 3.77.

### Supplementary Figure S7

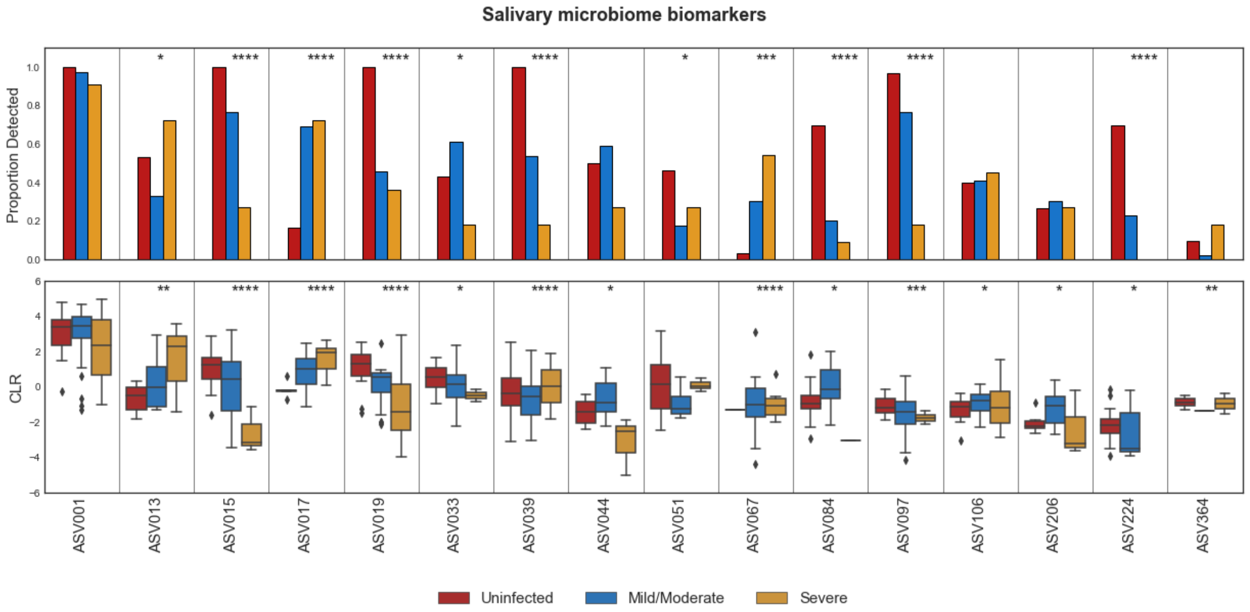


**Supplementary Figure S7.** Statistical significance for salivary microbiome biomarkers in the context of (top) detected vs. not detected using a Fisher's Exact test and (bottom) CLR values using a Kruskal-Wallis H-test. FDR < 0.05 *, FDR < 0.01 **, FDR < 0.001 ***, and FDR < 0.0001 ****.

### Supplementary tables

#### Supplementary table S1

| Sub-model | Modality | Features |
|---|---|---|
| **y1** | **Microbiome** | ASV017, ASV019, ASV033, ASV067, ASV001, ASV084, ASV039, ASV224, ASV051, ASV206 |
| | **Salivary** | IL16, ENA78CXCL5, VEGFA, IL17ACTLA8, IL5, TWEAK, FGF2, FractalkineCX3CL1, HGF, GROalphaCXCL1, MCP2CCL8, NGFbeta, MIP1alphaCCL3, MCP1CCL2, MIF, IL27, TNFalpha, MMP1, IL4, MCSF, TNFRII, MIGCXCL9, IP10CXCL10, IL1beta, GCSFCSF3, IL10 |
| | **Systemic** | MDC, IP10_CXCL10, HG, MCP2_CCL8, MIP1_btea_CCL4, MIP3_alpha_CCL20 |
| **y2** | **Microbiome** | ASV013, ASV015, ASV044, ASV106, ASV364, ASV097 |
| | **Salivary** | IL4, FGF2, VEGFA, IP10CXCL10, IL16, MMP1, TNFalpha, MIF, IL10, IL8CXCL8, IL6, IL5 |
| | **Systemic** | Eotaxin2_CCL24, MIG_CXCL9, MCSF, MCP3_CCL7, TWEAK, HG, MIP1_btea_CCL4, IL7, MCP1_CCL2, BAFF, Eotaxin3_CCL26, IL2R, IFN_alpha, APRIL, IL22 |

**Supplementary Table S1.** Biomarker descriptors for abundance-based paradigms with respect to sub-model

**Supplementary Table S2**

| ASV_ID | Phylum | Class | Order | Family | Genus |
|---|---|---|---|---|---|
| **ASV001** | Firmicutes | Negativicutes | Veillonellales-Selenomonadales | Veillonellaceae | Veillonella |
| **ASV013** | Firmicutes | Negativicutes | Veillonellales-Selenomonadales | Veillonellaceae | Veillonella |
| **ASV015** | Firmicutes | Bacilli | Staphylococcales | Gemellaceae | Gemella |
| **ASV017** | Firmicutes | Negativicutes | Veillonellales-Selenomonadales | Veillonellaceae | Veillonella |
| **ASV019** | Proteobacteria | Gammaproteobacteria | Pasteurellales | Pasteurellaceae | |
| **ASV033** | Actinobacteriota | Actinobacteria | Micrococcales | Micrococcaceae | Rothia |
| **ASV039** | Actinobacteriota | Actinobacteria | Corynebacteriales | Corynebacteriaceae | Corynebacterium |
| **ASV044** | Bacteroidota | Bacteroidia | Flavobacteriales | Flavobacteriaceae | Capnocytophaga |
| **ASV051** | Firmicutes | Negativicutes | Veillonellales-Selenomonadales | Veillonellaceae | Veillonella |
| **ASV067** | Cyanobacteria | Cyanobacteriia | Chloroplast | Chloroplast | Chloroplast |
| **ASV084** | Bacteroidota | Bacteroidia | Bacteroidales | Prevotellaceae | Alloprevotella |
| **ASV097** | Firmicutes | Bacilli | Erysipelotrichales | Erysipelotrichaceae | Solobacterium |
| **ASV106** | Fusobacteriota | Fusobacteriia | Fusobacteriales | Leptotrichiaceae | Leptotrichia |
| **ASV206** | Actinobacteriota | Actinobacteria | Bifidobacteriales | Bifidobacteriaceae | Alloscardovia |
| **ASV224** | Bacteroidota | Bacteroidia | Bacteroidales | Rikenellaceae | Rikenellaceae_RC9_gut_group |
| **ASV364** | Firmicutes | Negativicutes | Veillonellales-Selenomonadales | Selenomonadaceae | |

**Supplementary Table 2**. Taxonomic lineage for salivary microbiome biomarkers.
